# Modulations of thalamo-cortical coupling during voluntary movement in patients with essential tremor

**DOI:** 10.1101/2025.02.15.638416

**Authors:** Alexandra Steina, Sarah Sure, Markus Butz, Jan Vesper, Alfons Schnitzler, Jan Hirschmann

**Author notes:** Correspondence to: Jan Hirschmann, Full address: Moorenstraße 5, 40225 Düsseldorf, Germany.

## Abstract

The ventral intermediate nucleus of the thalamus (VIM) is the main thalamic hub for processing cerebellar inputs and the main deep brain stimulation target for the treatment of essential tremor (ET). As such, it presumably plays a critical role in motor control. So far, however, this structure has been rarely investigated in humans, and almost all of the existing studies focus on tremor. Here, we set out to study neural oscillations in the VIM and their coupling to cortical oscillations during voluntary movement.

We investigated thalamo-cortical coupling by means of simultaneous recordings of thalamic local field potentials and magnetoencephalography in 10 ET patients with externalized deep brain stimulation electrodes. Brain activity was measured while patients were pressing a button repeatedly in response to a visual cue. In a whole-brain analysis of coherence between VIM and cortex, we contrasted activity around a pre-movement baseline and button pressing.

Button pressing was associated with a bilateral decrease of thalamic power in the alpha (8– 12 Hz) and beta (13–21 Hz) band and a contralateral power increase in the gamma (35– 90 Hz) band. Moreover, changes in VIM-cortex coherence were observed. Alpha/low beta (8– 20 Hz) coherence decreased before and during movement, and the effect localized to the supplementary motor area and premotor cortex. A rebound of high beta (21–35 Hz) coherence occurred in the same region, but was more focal than the suppression. Pre-movement levels of thalamo-cortex low-beta coherence correlated with reaction time.

Our results demonstrate that voluntary movement is associated with modulations of behaviourally relevant thalamic coupling, primarily to premotor areas. We observed a clear distinction between low- and high-beta frequencies and our results suggest that the concept of “antikinetic” beta oscillations, originating from research on Parkinson’s disease, is transferable to ET.

## Introduction

The ventral intermediate nucleus of the motor thalamus (VIM) is believed to play a major role in the pathophysiology of essential tremor (ET).^1^ Deep brain stimulation (DBS) of the VIM effectively suppresses tremor and oscillatory activity in the VIM has been demonstrated to be coherent with activity from the tremulous limb during tremor.^2^

Apart from its role in tremor, the motor thalamus is involved in controlling voluntary movements, maintaining postures, and motor learning.^3,4^ Local field potential (LFP) recordings, for example, have revealed that oscillatory activity in the VIM is modulated during voluntary movements. During both self-paced and externally triggered movements, beta activity (13–30 Hz) decreases, while gamma activity (35–90 Hz) increases.^5–7^ Such movement-related modulations of oscillatory activity are a ubiquitous phenomenon, occurring in several motor-related brain areas, such as motor cortex^8^ or basal ganglia.^9^ Beta activity is often interpreted as antikinetic,^10,11^ i.e. anti-correlated with movement speed, while gamma activity is considered pro-kinetic.^12^

Expanding on these findings, studies combining LFP and cortical recordings have revealed that subcortical-cortical coupling follows similar dynamics. For example, in dystonia patients, low-beta (13–21 Hz) GPi-cortex coupling diminishes during cued movements, with coherence values correlating with reaction times,^11^ in line with an antikinetic nature of beta oscillations. Similarly, in Parkinson’s disease, movement onset is accompanied by suppression of beta coherence and an increase of gamma coherence between the STN and cortex,^13^ with levodopa-induced bradykinesia improvements correlating with greater gamma coherence.^9^ There is also initial evidence for modulations of thalamo-cortical coupling during voluntary movement,^6,7^ but their topography, dynamics, and behavioural relevance remain elusive.

The aim of this study was to describe the network topology of thalamo-cortical coupling and its dynamic modulations before, during, and after voluntary movement. For this purpose, we performed LFP recordings from externalized VIM-DBS electrodes in combination with whole-head MEG during externally triggered button pressing in patients with ET. Moreover, we correlated coherence values with reaction times to demonstrate the behavioural relevance of thalamo-cortical coupling.

## Materials and methods

### Patients and recordings

10 patients diagnosed with essential tremor, undergoing surgery for DBS, participated in the study. Before the recording, patients provided written informed consent according to the Declaration of Helsinki and the study was approved by the Ethics Committee of the Medical Faculty at Heinrich Heine University Düsseldorf (ET: “2018-217-Zweitvotum”). The measurements happened the day after implantation of DBS electrodes and before the pulse generator was implanted, allowing for the recording of LFPs from the externalized leads. Patient details are provided in **Table 1**.

**Table 1.** Patient details.

| Patient ID | age [y] | sex | disease duration [y] | electrode type |
| --- | --- | --- | --- | --- |
| ET01 | 65 | m | 19 | Abbott Infinity |
| ET02 | 71 | m | 20 | Abbott Infinity |
| ET03 | 60 | f | 49 | Abbott Infinity |
| ET04 | 62 | m | 50 | Abbott Infinity |
| ET05 | 57 | f | 51 | Abbott Infinity |
| ET06 | 76 | m | n.a. | Abbott Infinity |
| ET07 | 54 | m | 39 | Medtronic SenSight |
| ET08 | 62 | f | 56 | Abbott Infinity |
| ET09 | 65 | m | 20 | Boston Sc. Vercice Cartesia |
| ET10 | 71 | m | 20 | Boston Sc. Vercice Cartesia |
| $\mu \pm \sigma$ | $65 \pm 11$ | | $31 \pm 20$ | |

y: year, m: male, f: female, n.a.: not available, $\mu$ : mean, $\sigma$ : standard deviation.

### Electrophysiological measurements

We recorded MEG combined with intracranial LFPs from bilateral electrodes targeting the VIM. The LFPs were referenced to a mastoid reference. MEG signals were recorded by a 306-channel MEG system (Vectorview, MEGIN) with a sampling rate of 2 kHz. Moreover, we measured electromyograms (EMGs) from both forearms (extensor digitorum communis and flexor digitorum communis), accelerometer signals from both index fingers, and vertical and horizontal electrooculograms.

### Paradigm

The experiment included one motor task, which followed a resting state recording^14^ and two other motor tasks,^15^ that were analysed in our previous works.

During the motor task, a button box was placed on a table in front of the patients. Upon the presentation of a visual cue, presented on a screen in front of them, patients pressed a button with either the left or the right index finger. Each trial started with a black fixation cross that was presented between 6-8 s, followed by a Go cue (green cross). Pressing the button started the next trial. The task was performed in blocks of 8 min and each patient completed 1-3 blocks. Each block was divided equally into a left-hand and a right-hand part, and started with a short video indicating which index finger to use first. The hand was switched after half of the trails had been recorded, with the hand switch indicated by a second video.

### Data preprocessing

Data analysis was performed with the FieldTrip toolbox,^16^ MNE-Python,^17^ and custom written MATLAB (the MathWorks) scripts. Raw data were scanned for bad MEG and LFP channels and bad channels were excluded from further analyses. In order to reduce artefacts we applied temporal signal space separation to the MEG data using MNE-Python’s *mne*.*preprocessing*.*maxwell_filter*, with *st_duration* set to 10 s and *st_correlation* to 0.98.

All following analysis steps were performed with the FieldTrip toolbox. For further analysis, we used only the 204 planar gradiometers and down-sampled the data to 200 Hz. We rearranged the LFPs into a bipolar montage by subtraction of signals from adjacent electrode contacts. EMGs were high-pass filtered at 10 Hz and full-wave rectified.

### Epoching

The data were arranged in trials ranging from -4 to 4 s relative to button press (t = 0 s). Trials were visually inspected and bad trials were removed. Additional trials were discarded if the variance of any LFP channel exceeded 10^-8^ *µV* or if patients pressed the button more than once within the 8 s interval around the button press. One patient was excluded from further analysis because of bad LFP quality throughout the button pressing task. Information on the final number of trials can be found in **Supplementary Table 1**.

### Source reconstruction

We generated a single-shell head model for each patient based on the individual T1-weighted MRI scan (Siemens Mangetom Tim Trio, 3-T MRI scanner, München, Germany) and reconstructed sources for a grid with 567 points. The grid points were distributed over the cortical surface, aligned to Montreal Neurological Institute (MNI) space. For source reconstruction, we used a linear constrained minimum variance (LCMV) beamformer,^18^ with the regularization parameter *λ* set to 5%. Temporal signal space separation results in rank reduction, which can lead to erroneous beamformer output. To account for the rank reduction, we truncated the covariance matrix such that it had the same rank as the Maxwell-filtered data. To minimize confounds due to differences in spatial filters, we applied a common spatial filter to both condition contrasts (button press *vs*. baseline).

### Time-resolved spectra

For the trial-based data, we calculated time-resolved power and thalamo-cortical coherence spectra with a sliding window of 800 ms which was moved in steps of 50 ms. At each time step, complex Fourier spectra were calculated from 5-45 Hz and 55-90 Hz using multi-tapering with 2 Hz spectral smoothing, from which we derived power and coherence. The interval from 45-55 Hz was excluded due to 50 Hz line noise. For statistical analysis, we defined two intervals of interest: a baseline period from -3.0 to -2.0 s and a peri-movement interval from -1.5 to 2.5 s.

### Contact localization and contact selection

Using a pre-operative MRI and a post-operative CT scan, we localized DBS electrodes with Lead-DBS.^19^ The localized electrodes are displayed in **Fig. 1C**. We ensured that electrodes were on target and used only contacts within the ventral thalamus for further analysis. Moreover, we selected one bipolar LFP channel showing the strongest 8-20 Hz desynchronization contralateral to the button press. Because we alternated blocks of left- and right-hand button pressing, this procedure resulted in two selected channels per patient, i.e. one per hemisphere.

**Figure 1:**
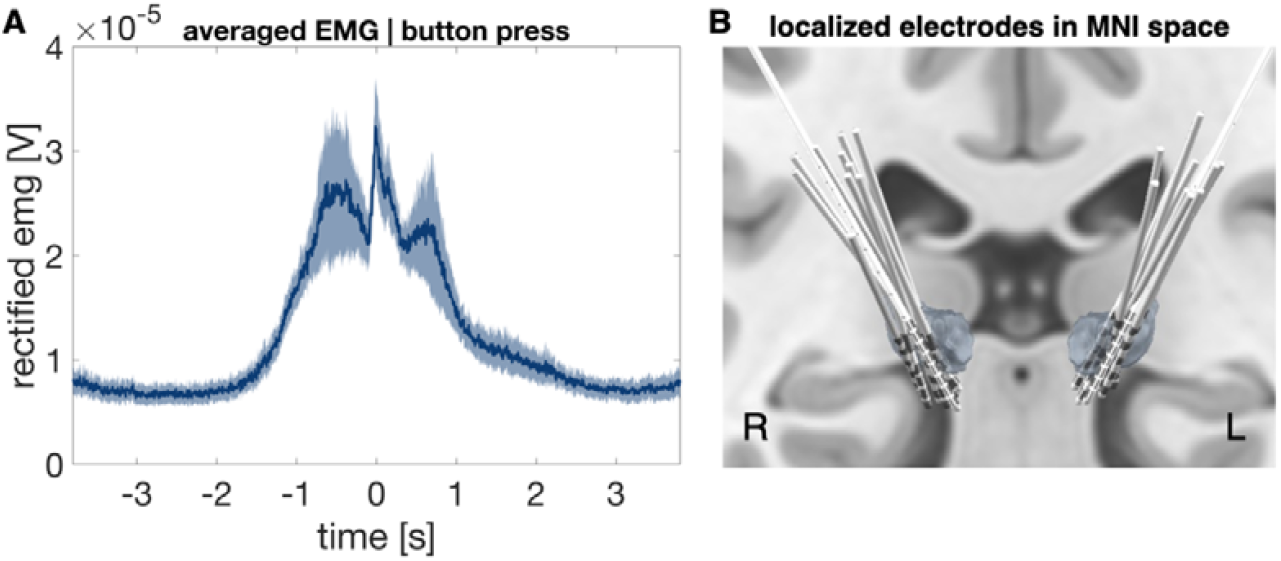
Electromyography signals and deep brain stimulation electrodes targeting the ventral intermediate nucleus of the thalamus. **(A)** EMG timeseries averaged over all patients, aligned to button press (t = 0). The shaded blue area represents the standard error of mean. **(B)** Electrodes targeting the VIM localized with Lead-DBS.

#### Source images

We computed band-limited coupling between reconstructed sources and LFPs for three frequency ranges of interest: alpha/low-beta (8-20 Hz), high beta (21-35 Hz), and gamma (65-85 Hz). Alpha and low-beta were aggregated as they changed jointly in the button press task. We applied bandwidth-wide spectral smoothing to capture an entire band in one estimate, using multi-tapering.^20^

For epochs containing right hand movement, we mirrored the source images across the midsagittal plane. In consequence, brain activity ipsilateral to movement appears in the left hemisphere in all figures, and brain activity contralateral to movement in the right hemisphere.

### Reaction time and pre-movement coherence

We tested if pre-movement coupling strength was predictive of reaction time. For this purpose, we calculated pre-movement coherence (-0.5 to 0.5 s relative to the Go cue) in the alpha/low-beta (8-20 Hz) and in the high beta (21-35 Hz) band and correlated it with reaction time. One patient’s reaction time was not stored due to technical problems.

### Statistical analysis

Rather than patient, the unit of observation of this study was hemisphere (*N*_*hemispheres*_ = 17) in line with previous studies.^14,21,22^ The statistical analysis had a within-hemisphere design (movement *vs*. baseline) and we used a nonparametric, two-sample, cluster-based-permutation tests with 1000 random permutations. The tests were two-tailed, with an *α*-level of 0.05. The results were corrected for multiple comparisons by relating all effects to the strongest effects observed in the permuted data (brain-wide or spectrum-wide).^23^ Cortical areas showing differences served as regions of interest for further analyses, such as Pearson correlation with behavioural metrics or visualization of power/coherence dynamics.

## Results

### Button pressing

In the button pressing task, patients had to press a button every 6-8 s in response to a visual cue. Group average EMG activity, aligned to the button press, is displayed in **Fig. 1B**. On average, movement started between 2-1.5 s before the button was pressed, as patients had to first lift their hand from the table and reach towards the button box.

### Movement-related power changes of thalamic oscillatory activity

Movement-related changes in VIM power ipsi- and contralateral to the button press are depicted in **Fig. 2**. VIM power in the 8–20 Hz range started to decrease below baseline levels ∼ 1 s before the button press, and this decrease lasted until ∼1.5 s after the button press. Besides this movement-related alpha/low-beta power suppression, which occurred bilaterally (ipsilateral VIM: cluster-based-permutation-test, *t*_*clustersum*_ = -1.61*10^3^, *p* = 0.004; contralateral VIM: *t* = –2.5*10^3^, *p* = 0.002), we observed movement-locked power increases with a pronounced hemispheric lateralization. In the VIM contralateral to movement, power in the 21-35 Hz range increased around 0-2 s relative to the button press (*t* = 1.7*10^3^, *p* = 0.003), likely reflecting a combination of a low-gamma power increase around movement onset and a post-movement beta rebound. A further gamma power increase was observed at higher frequencies (65-85 Hz), around the time of button press (*t* = 746, *p* < 0.001).

**Figure 2:**
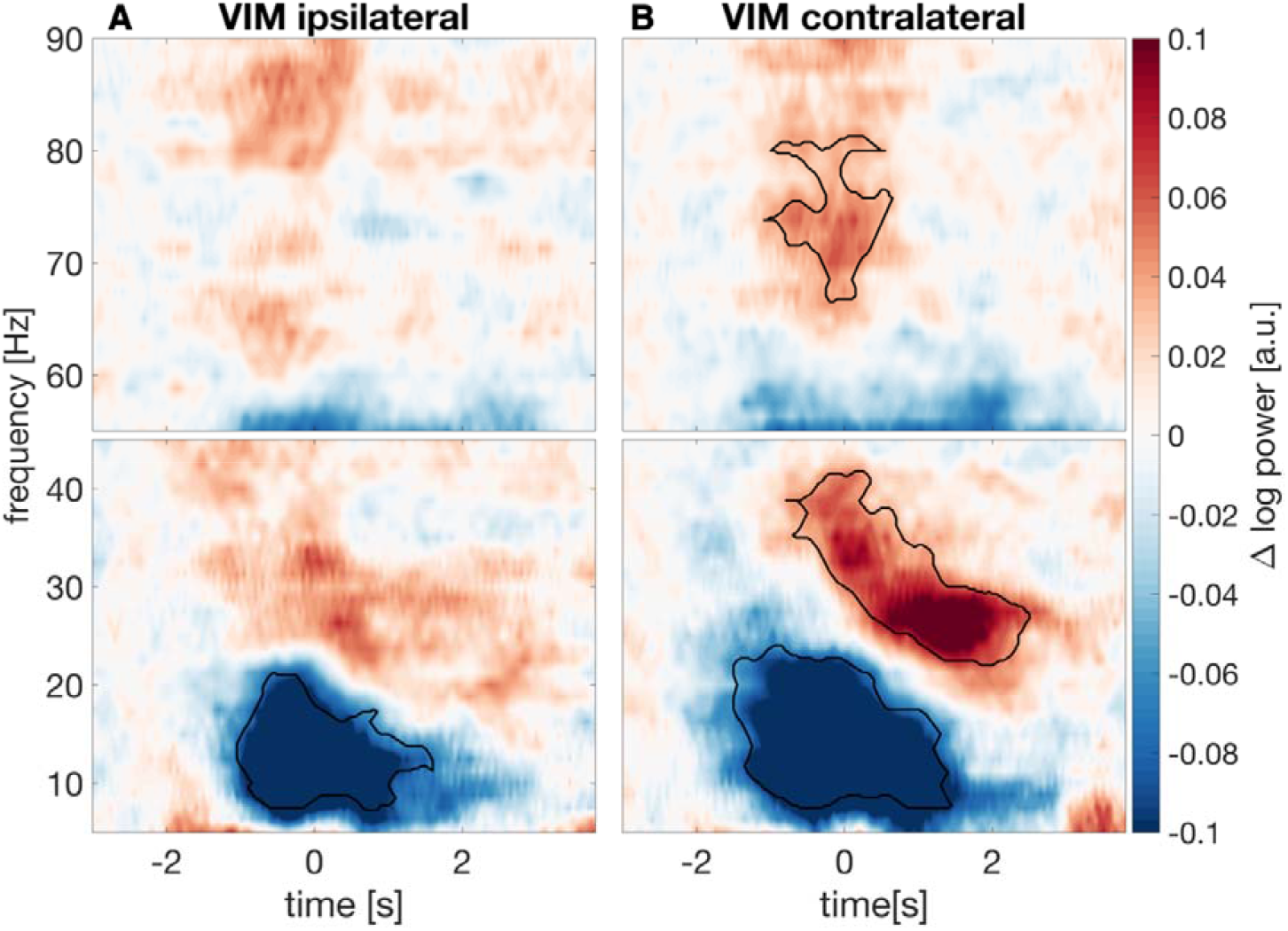
Thalamic power is modulated during button pressing. Baseline-corrected time frequency spectra of VIM power averaged over 17 hemispheres from nine patients in the hemisphere **(A)** ipsilateral and **(B)** contralateral to the button press (time point 0 s). Colours code absolute change in log10-transformed power compared to the mean baseline level (–3.0 – –2.0 s). Black contours mark significant changes (*p* < 0.05).

### Movement related changes of VIM-cortex coherence

For the whole-brain VIM-cortex coherence analysis, we defined three time-frequency intervals of interest: -0.5-0.5 s/8-20 Hz, -0.5-0.5 s/ 65-85 Hz and 0.5-1.5 s/21-35 Hz. These intervals reflect the movement-related alpha/low-beta power suppression, the gamma power increase locked to the button press and the high-beta power rebound, respectively, as observed for VIM power (**Fig. 2**). We assessed changes of VIM-cortex coherence in these intervals relative to baseline for the hemisphere contralateral to movement and for the hemisphere ipsilateral to movement.

### Contralateral hemisphere

For the first time-frequency interval of interest (-0.5-0.5 s/8-20 Hz; movement-related alpha/low-beta suppression), we observed a decrease of VIM-cortex coherence in primary motor, premotor, and primary somatosensory cortex contralateral to movement (cluster-based-permutation test; *t*_*clustersum*_ = –160, *p* = 0.002; MNI-coordinates minimal t-value: X = +/– 54.4 mm, Y = –40 mm, Z = 51.1 mm), which was strongest in superior frontal gyrus. For the second time-frequency interval of interest (0.5-1.5 s/21-35 Hz; high-beta rebound), we found an increase of coherence with precentral gyrus (*t* = 46.2, *p* = 0.003; X = +/–39.7 mm, Y = 0 mm, Z = 59.4 mm). This coherence rebound was mostly contained within the region presenting the movement-related suppression earlier in the trial (**Fig. 3**). Peri-movement VIM-cortex coherence changes in the gamma-band (-0.5-0.5 s/ 65-85 Hz) mapped to similar areas (**Supplementary Fig. 1**) but were not significant (*t* = 15.4, *p* = 0.09; X = 39.7 mm, Y = 0 mm, Z = 59.4 mm).

**Figure 3:**
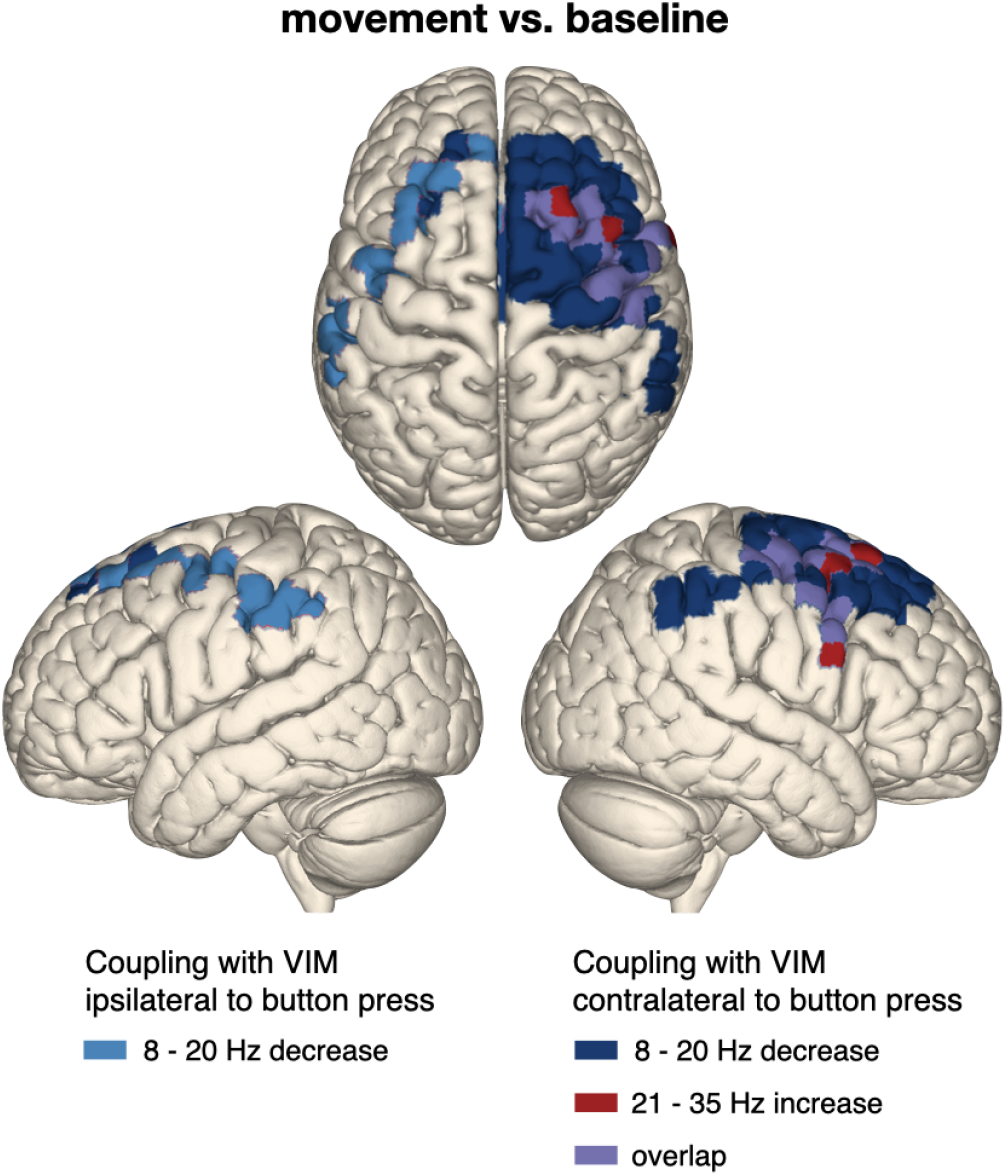
Thalamo-cortical coupling is modulated during button pressing. Coupling between cortex and VIM contralateral (dark blue) and ipsilateral (light blue) to button press decreased in the 8-20 Hz range during the button press, after which beta coherence rebounded in the 21-35 Hz range (red). The overlap between movement-related beta suppression and post-movement beta rebound is marked in purple. Non-significant changes are masked. Left hemisphere: ipsilateral to button press, right hemisphere: contralateral to button press. Note that the colours code significant effects rather than effect size.

### Ipsilateral hemisphere

For the VIM ipsilateral to movement, we observed a movement-related alpha/low-beta suppression, which mapped to the supramarginal gyrus ipsilateral to button press (*t* = –38.8, *p* = 0.012; X = –/+56.4 mm, Y = –30 mm, Z = 46.4 mm; **Fig. 3**), but no significant rebound.

### Dynamics of thalamo-cortical coupling

To investigate the dynamics of thalamo-cortical coupling during button pressing, we computed a time-frequency spectrum of coherence between the VIM and the region with the strongest changes in the whole-brain analysis (bilateral motor and pre-motor cortex; see **Fig. 3**). Because the former analysis had already revealed significant deviations from baseline, we did not re-assess significance here.

The dynamics of coherence resembled those of VIM power (**Fig. 2**), with a peri-movement alpha/low-beta suppression in both the contra- and the ipsilateral hemisphere (with respect to movement). The post-movement beta rebound in the high-beta range was stronger in the hemisphere contralateral to movement (**Fig. 4**).

**Figure 4:**
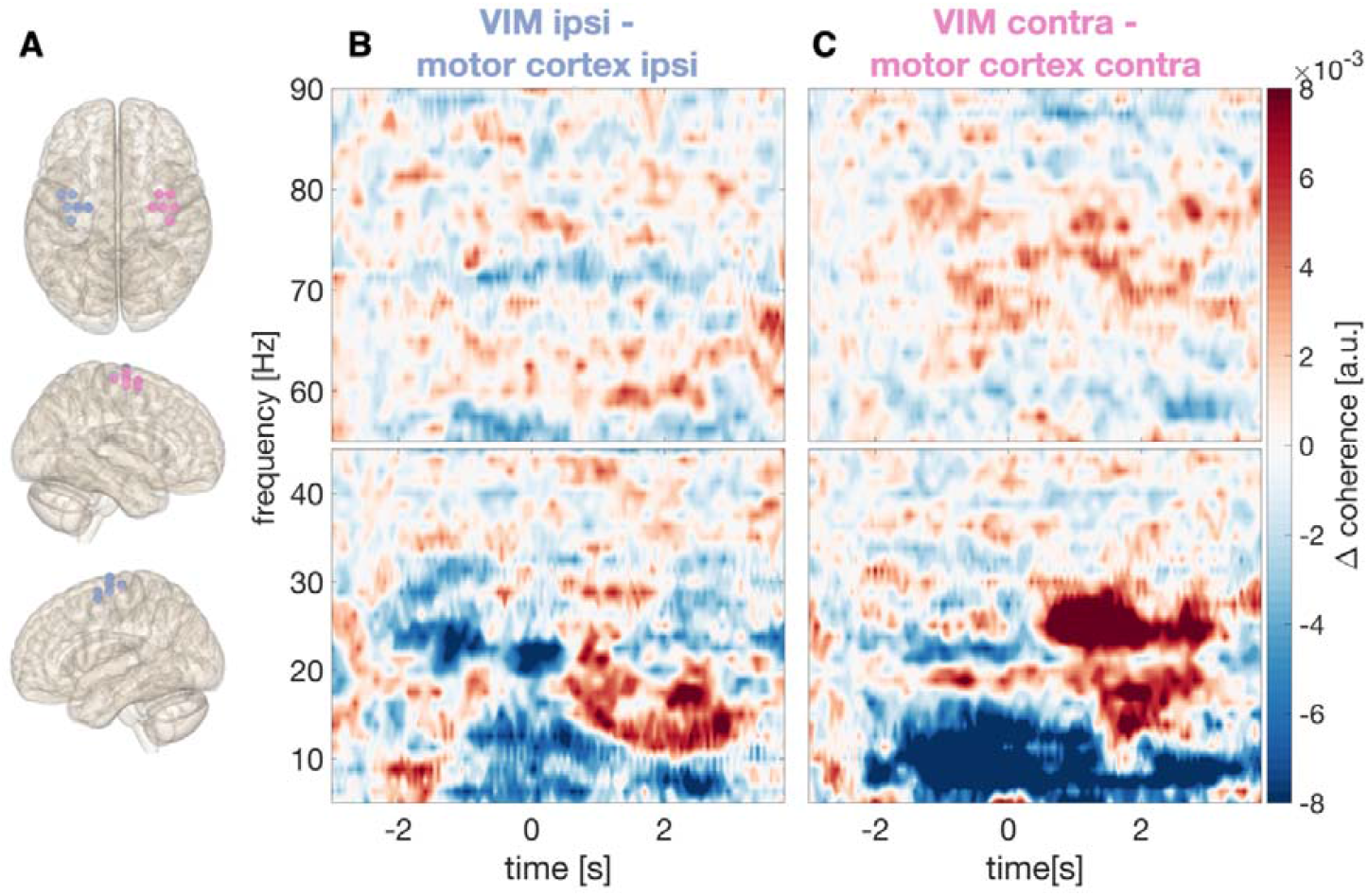
Time-resolved dynamics of thalamo-cortex coherence during button pressing. **(A)** Grid points (beamformer target locations) defining the regions of interest. Coherence was computed for each grid point and averaged within the region of interest. Left hemisphere: ipsilateral to button press, right hemisphere: contralateral to button press. (**B**-**C**) Baseline-corrected time frequency spectra of coherence between VIM and motor cortex **(B)** ipsilateral and **(C)** contralateral to button press (time point 0 s).

### Dynamics of motor cortical power

We selected the same regions of interest as for coherence and computed time-resolved power spectra for the motor cortex ipsi- and contralateral to movement (**Fig. 5**). A strong power suppression ranging from 5–35 Hz was visible in both hemispheres (ipsilateral motor cortex: cluster-based-permutation-test, *t*_*clustersum*_ = -7.9*10^3^, *p* < 0.001; contralateral motor cortex: *t* = –1.1*10^4^, *p* < 0.001). We did neither observe a strong beta rebound, nor a gamma increase in motor cortex. To examine whether beta power was rebounding in motor cortex, we aligned the time-resolved power spectra to the time point when the button was released (**Supplementary Figure 2**). This analysis demonstrated a weak rebound in the alpha/beta range in the hemisphere contralateral to movement.

**Figure 5:**
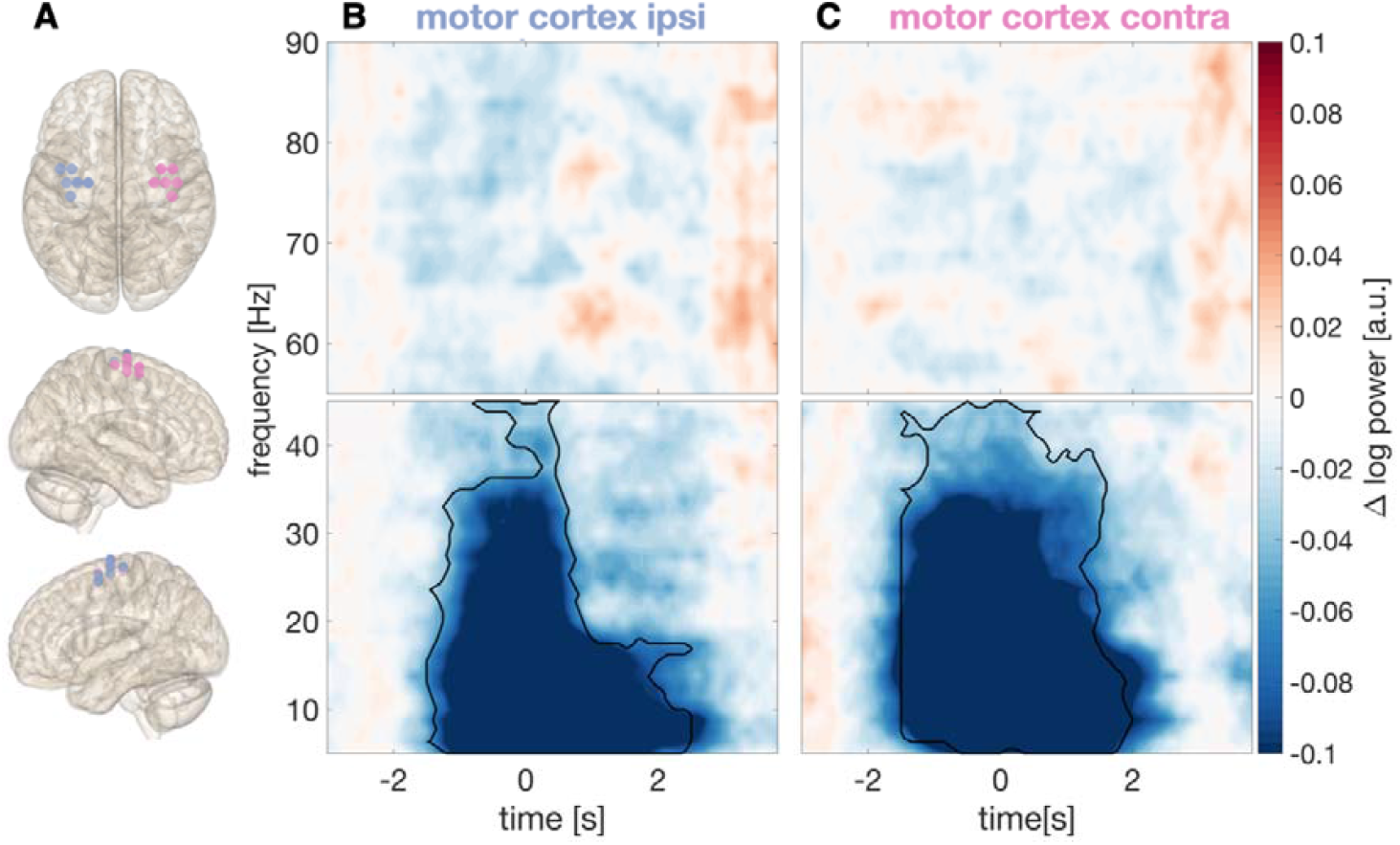
Time-resolved dynamics of cortical power during button pressing. **(A)** Grid points (beamformer target locations) defining the regions of interest. Left hemisphere: ipsilateral to button press, right hemisphere: contralateral to button press. (**B**-**C**) Baseline-corrected time frequency spectra of cortical power **(B)** ipsilateral and **(C)** contralateral to the button press (time point 0 s).

### Pre-movement VIM-cortex coherence and reaction time

Based on studies of Parkinson’s disease, beta band synchronisation has been labelled “antikinetic”, i.e. inversely related to movement speed. Here, we tested the validity of this label in ET patients by correlating pre-movement levels of VIM-motor cortex beta coherence (8–20 Hz, 21–35 Hz; - 0.5–0.5 s around Go cue) to reaction time (time of button press - time of Go cue presentation).

Pre-movement levels of 8–20 Hz coupling between VIM and motor cortex contralateral to movement was positively correlated with reaction time (*r* = 0.53, *p* = 0.038; **Fig. 6B**). The correlation was not significant for the high-beta band (*r* = -0.06, *p* = 0.82). Coupling between VIM and motor cortex ipsilateral to button press was not significantly correlated with reaction time, neither in the alpha/low-beta (8–20 Hz: *r* = 0.09, *p* = 0.74) nor in the high-beta band (**Fig. 6B;** 21–35 Hz: *r* = -0.45, *p* = 0.09).

**Figure 6:**
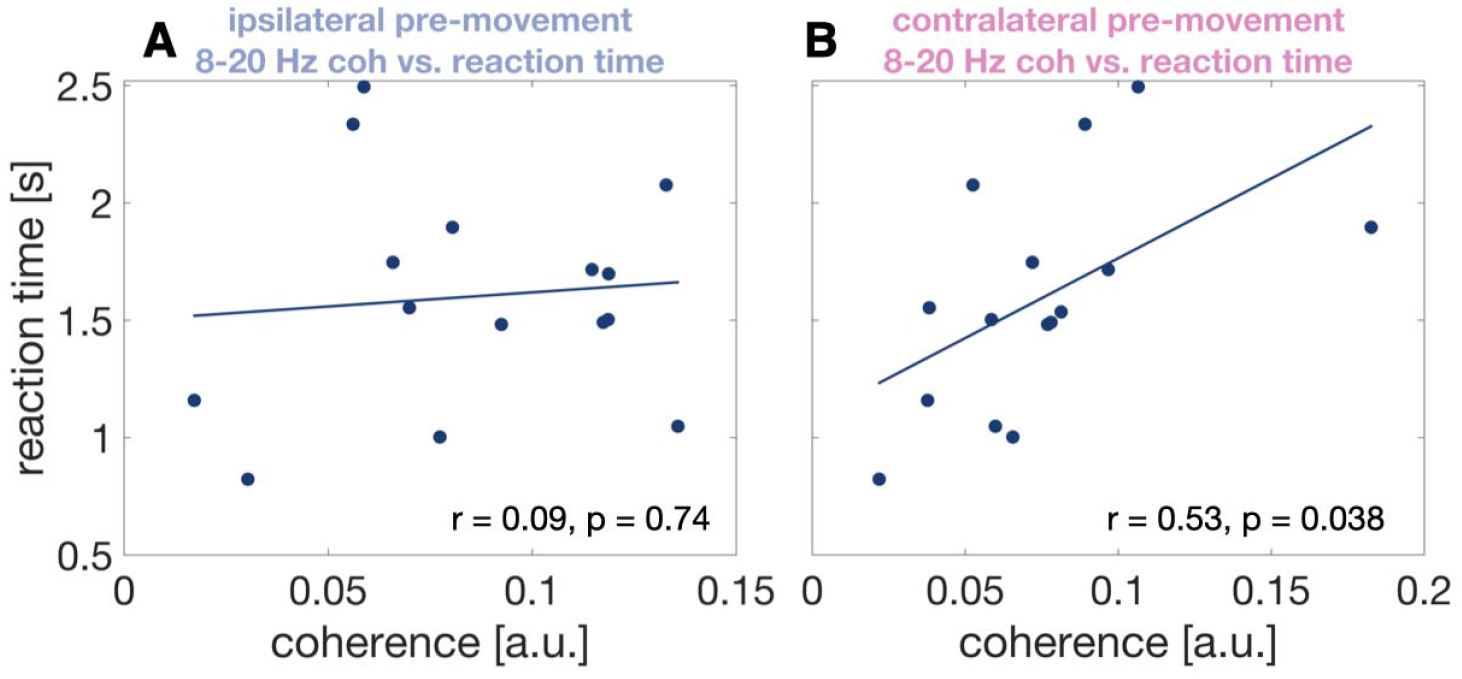
Correlation between coherence and reaction time. Scatterplot illustrating the relationship between pre-movement alpha/low-beta coherence and reaction time for coupling between ipsilateral VIM and ipsilateral motor cortex **(A)** and contralateral VIM and contralateral motor cortex **(B)**.

## Discussion

In the present study, we revealed the brain areas and frequency bands involved in thalamo-cortical coupling during voluntary movements. We found that voluntary movement is associated with peri-movement modulation of beta band coherence, involving mostly supplementary motor area and premotor cortex. Pre-movement alpha/low-beta coherence between motor cortex and VIM contralateral to movement correlated with reaction time, suggesting that beta band synchronization is generally associated with slowness, even in the absence of akinesia.

Our study is one of few works relating thalamo-cortical coupling to voluntary movement. Most studies have investigated tremor, which has a different coupling profile, involving other frequency bands and other brain areas.^24–26^ In fact, we have recently described tremor-related coherence profiles of half the patients analyzed in this work.^15^ Strongest tremor-associated modulations of VIM-cortex coherence were observed in primary sensorimotor cortex rather than pre-motor areas, suggesting that different channels of thalamo-cortical communication might be active during tremor and voluntary movements. A further difference might be the modulation of VIM-cerebellar coupling, which was pronounced for tremor but non-significant for button pressing. However, voluntary movement and tremor seem to share some common frequency-specific modulations. Voluntary movement is linked to a suppression in the beta-band, for example, and tremor amplitude is also inversely related to beta-band VIM-motor cortex coupling.^15^

Overall, our results underscore how closely neuronal oscillations in the motor system are linked to the movement present at the time of recording – a link which should be kept in mind when attributing oscillatory patterns to any specific disease.

### VIM power

Here, we reproduced previous findings on modulations of thalamic activity during voluntary movements.^5,7^ Before and while the button was pressed, alpha (8-12 Hz) and low-beta (13-20 Hz) activity were desynchronized, while gamma activity was synchronized. Shortly after the button press, high-beta (21-35 Hz) oscillations increased to a higher level than baseline (rebound). This pattern has been replicated in numerous motor structures, such as motor cortex^27^ or the STN,^9^ and numerous cohorts, including patients with Parkinson’s disease,^9^ dystonia,^11^ and healthy controls.^8^ Matching such a ubiquitous motive, we suggest that the spectral modulations of VIM activity observed here are of physiological rather than of pathophysiological nature.

The apparent divergence between the dynamics of low- and high-beta activity matches findings in Parkinson’s disease indicating distinct roles of low- and high-beta activity.^28,29^ Whether alpha oscillations change independently of low-beta oscillations or result from spectral leakage from the beta band is under debate.^30,31^ Whereas Klostermann et al. suggested a distinction between alpha and beta (13-35 Hz) activity,^7^ we found no evidence for independence between thalamic alpha and low-beta oscillations and treated both bands as a single entity.

### Motor cortical power

Motor cortical power largely resembled VIM power, but, in contrast to the VIM, motor cortex did not reveal a strong beta rebound in this paradigm. However, a weak rebound was visible in the hemisphere contralateral to movement when the spectral modulations were related to the time when the button was released (**Supplementary Fig. 2**). This finding suggests a differential response of the thalamus and motor cortex to single elements of the motor sequence.

### VIM-cortex coupling

VIM-cortex coupling followed a similar pattern as VIM power: alpha-/low-beta coherence decreased prior to and throughout the button press, and high-beta coherence increased after the button press. The decrease of alpha-/low-beta coherence has been reported before for externally paced^7,32^ and self-paced movements.^6^ We extend previous findings by localizing the coherence decrease to premotor cortex and the supplementary motor area. This localization is different from the spatial minimum of the cortical beta power suppression, which is typically observed in sensorimotor cortex proper, around the hand knob.^8^

Compared to baseline, high-beta coupling between cortex and the VIM contralateral to movement increased shortly after the button press. This effect was strongest in similar regions as the preceding 8–20 Hz decrease, but more focal. The coherence increase might be analogous to the post-movement rebound of beta power that is typically observed in sensorimotor cortex.^8^ This rebound is usually lateralized to the contralateral hemisphere,^8^ while the beta power suppression has been reported to be bilaterally symmetric.^33,34^ This pattern is in line with our results, as coherence decreased bilaterally during movement, whereas the post-movement coherence increase was lateralized to the contralateral hemisphere.

### VIM-cortex coupling and reaction time

Pre-movement levels of 8-20 Hz VIM-motor cortex coherence in the contralateral hemisphere correlated positively with reaction time, i.e. higher coupling around Go cue onset were associated with slower responses. These findings tally with the proposed antikinetic nature of low-beta oscillations, derived mainly from studies on Parkinson’s disease. These studies have established a relationship between elevated beta power in the STN and the severity of bradykinesia and rigidity.^35,36^ Moreover, movements of healthy individuals have been demonstrated to be slower when low-beta activity happens to be elevated in the course of spontaneous fluctuations or is elevated artificially by transcranial alternating current stimulation.^37,38^ Further, in patients with Parkinson’s disease deep brain stimulation in the beta range has shown to slow movements.^39^ The antikinetic nature of low-beta oscillations is not only reflected by local oscillatory power, but extends to coupling between different regions, as observed for GPi-cortex^11^ and cortico-spinal coherence,^40^ for example. Here, we demonstrate that the concept holds for VIM-cortex coupling, too.

Interestingly, other pre-movement features of thalamic activity have likewise been linked to reaction time. The amplitude of the contingent negative variation in between a pre- and a Gocue was predictive of reaction time in a cued Go/NoGo task.^41^ Moreover, increased levels of thalamic gamma activity have been revealed to result in faster task performance.^5^ These observations align well with the modern view on the role of the thalamus in motor control. For a long time, the motor thalamus was believed to simply relay inputs from cerebellum to motor cortex. However, in the last decades, it became evident that information to cortex is not just passively relayed, but modified by the thalamus.^3^ Our study further supports this notion.

### Limitations

Intracranial recordings from the human thalamus are only possible in patients. Therefore, we cannot be sure whether the oscillatory dynamics described here indeed relate to normal motor control. Yet, several of our findings match observations made in other patient populations and in healthy participants, who, at the cortical level, exhibit the same beta and gamma power dynamics at movement start and stop.^42^

Although the patient cohort consisted of individuals with ET, tremor was only present in 6 out of 17 analyzed body sides during button pressing most likely due to the stun effect (see **Supplementary Table 1**). Thus, we could not compute a meaningful statistical contrast between button pressing with intention tremor to button pressing without intention tremor, which would have been an interesting addition to our recent work on essential tremor .^15^

## Conclusions

Our study demonstrates behaviourally relevant modulations of thalamo-cortical coupling during voluntary movement. Further it extends the notion of beta oscillations being “antikinetic” to thalamo-cortical coupling.

## Data availability

Data can be made available in anonymized form upon reasonable request, conditional on patient consent.

## References

1. Neudorfer C, Kultas-Ilinsky K, Ilinsky I, et al. The role of the motor thalamus in deep brain stimulation for essential tremor. Neurotherapeutics. 2024;21(3):e00313. doi:10.1016/j.neurot.2023.e00313

2. Hua SE, Lenz FA. Posture-Related Oscillations in Human Cerebellar Thalamus in Essential Tremor Are Enabled by Voluntary Motor Circuits. Journal of Neurophysiology. 2005;93(1):117–127. doi:10.1152/jn.00527.2004

3. Sommer MA. The role of the thalamus in motor control. Current Opinion in Neurobiology. 2003;13(6):663–670. doi:10.1016/j.conb.2003.10.014

4. Bosch-Bouju C, Hyland BI, Parr-Brownlie LC. Motor thalamus integration of cortical, cerebellar and basal ganglia information: implications for normal and parkinsonian conditions. Front Comput Neurosci. 2013;7. doi:10.3389/fncom.2013.00163

5. Brücke C, Bock A, Huebl J, et al. Thalamic gamma oscillations correlate with reaction time in a Go/noGo task in patients with essential tremor. NeuroImage. 2013;75:36–45. doi:10.1016/j.neuroimage.2013.02.038

6. Paradiso G. Involvement of human thalamus in the preparation of self-paced movement. Brain. 2004;127(12):2717–2731. doi:10.1093/brain/awh288

7. Klostermann F, Nikulin VV, Kühn AA, et al. Task-related differential dynamics of EEG alpha- and beta-band synchronization in cortico-basal motor structures: EEG oscillations in motor structures. European Journal of Neuroscience. 2007;25(5):1604–1615. doi:10.1111/j.1460-9568.2007.05417.x

8. Jurkiewicz MT, Gaetz WC, Bostan AC, Cheyne D. Post-movement beta rebound is generated in motor cortex: Evidence from neuromagnetic recordings. NeuroImage. 2006;32(3):1281–1289. doi:10.1016/j.neuroimage.2006.06.005

9. Litvak V, Eusebio A, Jha A, et al. Movement-Related Changes in Local and Long-Range Synchronization in Parkinson’s Disease Revealed by Simultaneous Magnetoencephalography and Intracranial Recordings. J Neurosci. 2012;32(31):10541–10553. doi:10.1523/JNEUROSCI.0767-12.2012

10. Engel AK, Fries P. Beta-band oscillations — signalling the status quo? Current Opinion in Neurobiology. 2010;20(2):156–165. doi:10.1016/j.conb.2010.02.015

11. van Wijk BCM, Neumann WJ, Schneider GH, Sander TH, Litvak V, Kühn AA. Low-beta cortico-pallidal coherence decreases during movement and correlates with overall reaction time. NeuroImage. 2017;159:1–8. doi:10.1016/j.neuroimage.2017.07.024

12. Brown P. Oscillatory nature of human basal ganglia activity: Relationship to the pathophysiology of Parkinson’s disease. Movement Disorders. 2003;18(4):357–363. doi:10.1002/mds.10358

13. Alegre M, Rodríguez-Oroz MC, Valencia M, et al. Changes in subthalamic activity during movement observation in Parkinson’s disease: Is the mirror system mirrored in the basal ganglia? Clinical Neurophysiology. 2010;121(3):414–425. doi:10.1016/j.clinph.2009.11.013

14. Steina A, Sure S, Butz M, Vesper J, Schnitzler A, Hirschmann J. Mapping Subcortico-Cortical Coupling—A Comparison of Thalamic and Subthalamic Oscillations. Movement Disorders. 2024;39(4):684–693. doi:10.1002/mds.29730

15. Steina A, Sure S, Butz M, Vesper J, Schnitzler A, Hirschmann J. Oscillatory coupling between thalamus, cerebellum and motor cortex in essential tremor. Published online November 11, 2024:2024.11.11.622917. doi:10.1101/2024.11.11.622917

16. Oostenveld R, Fries P, Maris E, Schoffelen JM. FieldTrip: open source software for advanced analysis of MEG, EEG, and invasive electrophysiological data. Computational intelligence and neuroscience. 2011;2011:1–9.

17. Gramfort A. MEG and EEG data analysis with MNE-Python. Front Neurosci. 2013;7. doi:10.3389/fnins.2013.00267

18. Van Veen BD, Van Drongelen W, Yuchtman M, Suzuki A. Localization of brain electrical activity via linearly constrained minimum variance spatial filtering. IEEE Transactions on biomedical engineering. 1997;44(9):867–880.

19. Horn A, Li N, Dembek TA, et al. Lead-DBS v2: Towards a comprehensive pipeline for deep brain stimulation imaging. NeuroImage. 2019;184:293–316. doi:10.1016/j.neuroimage.2018.08.068

20. Thomson DJ. Spectrum estimation and harmonic analysis. Proceedings of the IEEE. 1982;70(9):1055–1096. doi:10.1109/PROC.1982.12433

21. Oswal A, Cao C, Yeh CH, et al. Neural signatures of hyperdirect pathway activity in Parkinson’s disease. Nat Commun. 2021;12(1):5185. doi:10.1038/s41467-021-25366-0

22. Hirschmann J, Özkurt TE, Butz M, et al. Distinct oscillatory STN-cortical loops revealed by simultaneous MEG and local field potential recordings in patients with Parkinson’s disease. NeuroImage. 2011;55(3):1159–1168. doi:10.1016/j.neuroimage.2010.11.063

23. Maris E, Oostenveld R. Nonparametric statistical testing of EEG- and MEG-data. Journal of Neuroscience Methods. 2007;164(1):177–190. doi:10.1016/j.jneumeth.2007.03.024

24. Schnitzler A, Münks C, Butz M, Timmermann L, Gross J. Synchronized brain network associated with essential tremor as revealed by magnetoencephalography. Mov Disord. 2009;24(11):1629–1635. doi:10.1002/mds.22633

25. He F, Sarrigiannis PG, Billings SA, et al. Nonlinear interactions in the thalamocortical loop in essential tremor: A model-based frequency domain analysis. Neuroscience. 2016;324:377–389. doi:10.1016/j.neuroscience.2016.03.028

26. Marsden JF. Coherence between cerebellar thalamus, cortex and muscle in man: Cerebellar thalamus interactions. Brain. 2000;123(7):1459–1470. doi:10.1093/brain/123.7.1459

27. Salmelin R, Hámáaláinen M, Kajola M, Hari R. Functional Segregation of Movement-Related Rhythmic Activity in the Human Brain. NeuroImage. 1995;2(4):237–243. doi:10.1006/nimg.1995.1031

28. van Wijk BC, Beudel M, Jha A, et al. Subthalamic nucleus phase–amplitude coupling correlates with motor impairment in Parkinson’s disease. Clinical Neurophysiology. 2016;127(4):2010–2019.

29. Cao C, Litvak V, Zhan S, et al. Low-beta versus high-beta band cortico-subcortical coherence in movement inhibition and expectation. Neurobiology of Disease. 2024;201:106689. doi:10.1016/j.nbd.2024.106689

30. Brown P, Williams D. Basal ganglia local field potential activity: Character and functional significance in the human. Clinical Neurophysiology. 2005;116(11):2510–2519. doi:10.1016/j.clinph.2005.05.009

31. Khawaldeh S, Tinkhauser G, Shah SA, et al. Subthalamic nucleus activity dynamics and limb movement prediction in Parkinson’s disease. Brain. 2020;143(2):582–596. doi:10.1093/brain/awz417

32. Opri E, Cernera S, Okun MS, Foote KD, Gunduz A. The Functional Role of Thalamocortical Coupling in the Human Motor Network. J Neurosci. 2019;39(41):8124–8134. doi:10.1523/JNEUROSCI.1153-19.2019

33. Pfurtscheller G, Berghold A. Patterns of cortical activation during planning of voluntary movement. Electroencephalography and Clinical Neurophysiology. 1989;72(3):250–258. doi:10.1016/0013-4694(89)90250-2

34. Winkler L, Butz M, Sharma A, et al. Beta Waves in Action: Context-Dependent Modulations of Subthalamo-Cortical Synchronization during Rapid Reversals of Movement Direction. eLife. 2024;13. doi:10.7554/eLife.101769.1

35. Kühn AA, Kupsch A, Schneider GH, Brown P. Reduction in subthalamic 8–35 Hz oscillatory activity correlates with clinical improvement in Parkinson’s disease. European Journal of Neuroscience. 2006;23(7):1956–1960.

36. Neumann WJ, Degen K, Schneider GH, et al. Subthalamic synchronized oscillatory activity correlates with motor impairment in patients with Parkinson’s disease. Movement Disorders. 2016;31(11):1748–1751.

37. Gilbertson T, Lalo E, Doyle L, Lazzaro VD, Cioni B, Brown P. Existing Motor State Is Favored at the Expense of New Movement during 13-35 Hz Oscillatory Synchrony in the Human Corticospinal System. J Neurosci. 2005;25(34):7771–7779. doi:10.1523/JNEUROSCI.1762-05.2005

38. Pogosyan A, Gaynor LD, Eusebio A, Brown P. Boosting Cortical Activity at Beta-Band Frequencies Slows Movement in Humans. Current Biology. 2009;19(19):1637–1641. doi:10.1016/j.cub.2009.07.074

39. Werner LM, Schnitzler A, Hirschmann J. Subthalamic nucleus deep brain stimulation in the beta frequency range boosts cortical beta oscillations and slows down movement. J Neurosci. Published online January 7, 2025. doi:10.1523/JNEUROSCI.1366-24.2024

40. van Wijk BCM, Daffertshofer A, Roach N, Praamstra P. A Role of Beta Oscillatory Synchrony in Biasing Response Competition? Cerebral Cortex. 2009;19(6):1294–1302. doi:10.1093/cercor/bhn174

41. Nikulin VV, Marzinzik F, Wahl M, et al. Anticipatory activity in the human thalamus is predictive of reaction times. Neuroscience. 2008;155(4):1275–1283. doi:10.1016/j.neuroscience.2008.07.005

42. Alegre M, Gurtubay IG, Labarga A, Iriarte J, Valencia M, Artieda J. Frontal and central oscillatory changes related to different aspects of the motor process: a study in go/no-go paradigms. Exp Brain Res. 2004;159(1):14–22. doi:10.1007/s00221-004-1928-8

