## Supplementary for "Modulations of thalamo-cortical coupling during voluntary movement in patients with essential tremor"

***Supplementary material***

Alexandra Steina^1^, Sarah Sure^1^, Markus Butz^1^,
Jan Vesper^2^, Alfons Schnitzler^1^, Jan Hirschmann^1^

**Author affiliations:**

1 Institute of Clinical Neuroscience and Medical Psychology, Medical Faculty,
Heinrich Heine University, 40225, Düsseldorf, Germany

2 Department of Functional Neurosurgery and Stereotaxy, Neurosurgical Clinic,
Medical Faculty, Heinrich Heine University, 40225, Düsseldorf, Germany

**Supplementary Table 1 Task information**

L: left, R: Right, n.p.: not present, -.: not available, y: yes.

^a^Data length, tremor.

^b^Amount of trials after cleaning.

^c^Right VIM excluded due to uncertain electrode position.

^d^Button press from ET09 was excluded from further analysis due to bad LFP data quality.

| **Patient ID** | **Button press L / R tremor** | **Button press L / R trials^b^** |
| --- | --- | --- |
| ET01 | n.p. | 53 / 45 |
| ET02 | y / y | 83 / 85 |
| ET03 | n.p. | 89 / 87 |
| ET04 | n.p. / n.p. | 76 / 76 |
| ET05 | y / n.p. | 44 / 37 |
| ET06^c^ | y / n.p. | 54 / 54 |
| ET07 | n.p. | 63 / 59 |
| ET08 | y / y | 54 / 49 |
| ET09^d^ | y / y | - |
| ET10 | n.p. / y | 51 / 63 |

**Movement related changes of VIM-cortex coherence**

VIM-cortex coherence in the gamma range increased during movement. This increase was strongest in the pre-central gyrus, but the increase was not significant (*t_clustersum_* = 15.4, *p* = 0.09; X = 39.7 mm, Y = 0 mm, Z = 59.4 mm; **Supplementary Fig. 1**).

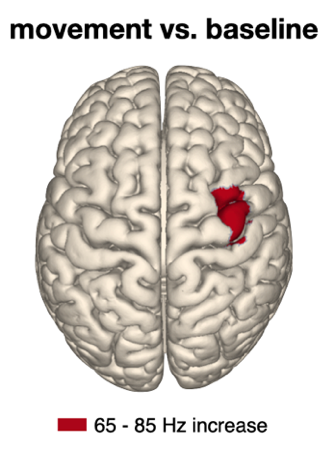

**Supplementary Fig. 1: Change of thalamo-cortical coupling in the gamma range during button pressing.** Coupling between cortex and the VIM contralateral to movement in the 65-85 Hz range increased during button pressing. Strongest changes are displayed (*t_clustersum_* = 15.4, *p* = 0.09; non-significant).

**Dynamics of motor cortical power**

We aligned time-resolved power in motor cortex to the time point when the button was released (t = 0) to investigate if beta power was rebounding in motor cortex. On a descriptive level, this analysis suggests a weak rebound in the alpha/beta range in the hemisphere contralateral to movement (**Supplementary Figure 2**). The weakness of the rebound might be due to the fact that the hand was still in motion after the button press, as patients returned their hand to the table in front of them. Note, however, the VIM and VIM-cortex coupling showed a marked rebound even in this situation.

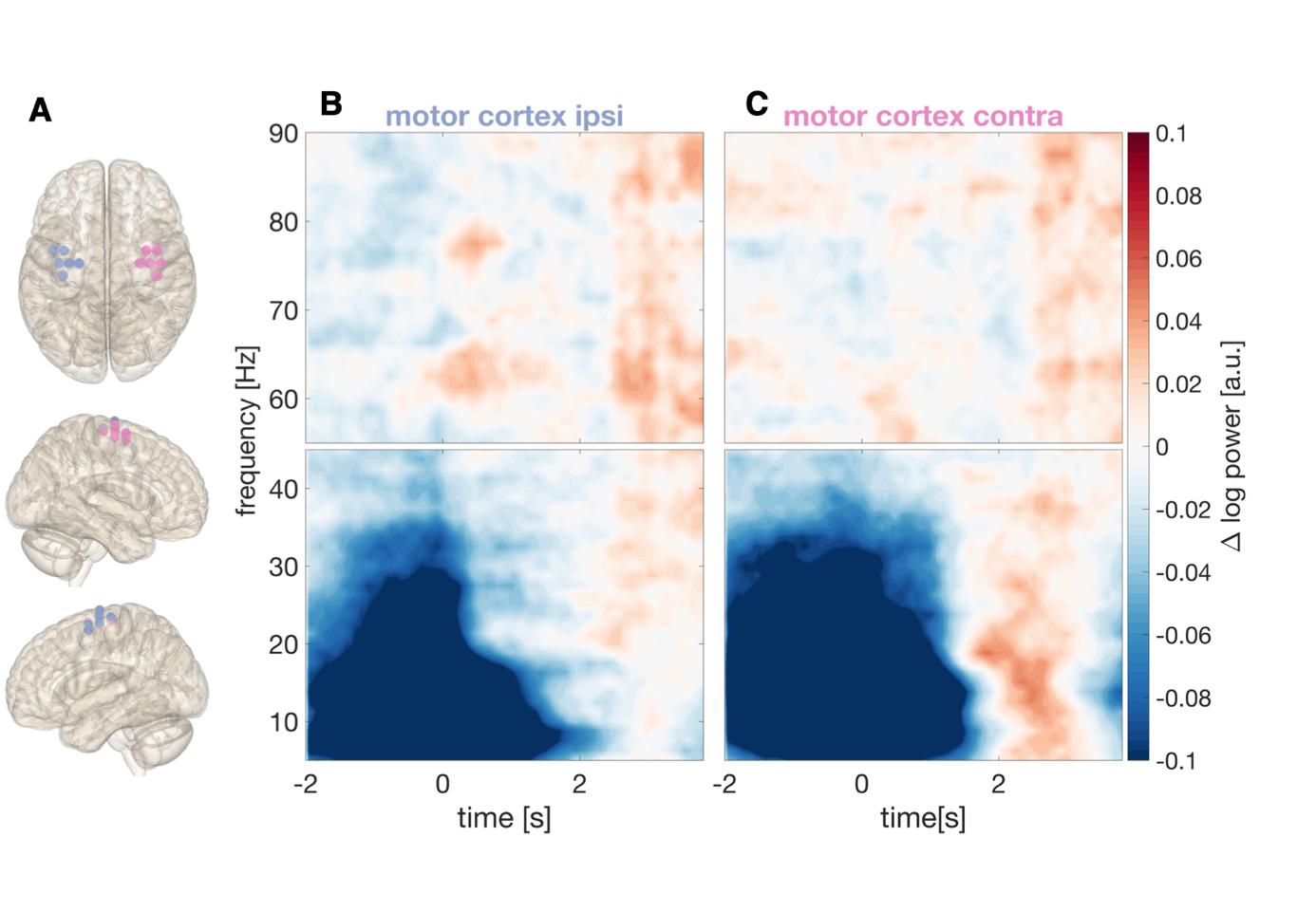

**Supplementary Fig. 2: Time-resolved dynamics of cortical power during button pressing aligned to end of button press. (A)** Grid points used for extracting cortical activity. Left hemisphere: ipsilateral to button press, right hemisphere: contralateral to button press. (**B**-**C**) Baseline-corrected (-3 to -2 s before onset of button press) time frequency spectra of cortical power **(B)** ipsilateral and **(C)** contralateral to the button press (time point 0 s marks the time when the button was released).
